# Rational design of a compact CRISPR-Cas9 activator for AAV-mediated delivery

**DOI:** 10.1101/298620

**Authors:** Suhani Vora, Jenny Cheng, Ru Xiao, Nathan J. VanDusen, Luis Quintino, William T. Pu, Luk H. Vandenberghe, Alejandro Chavez, George Church

## Abstract

Akin to Zinc Finger and Transcription Activator Like Effector based transcriptional modulators, nuclease-null CRISPR-Cas9 provides a groundbreaking programmable DNA binding platform, begetting an arsenal of targetable regulators for transcriptional and epigenetic perturbation, by either directly tethering, or recruiting, transcription enhancing effectors to either component of the Cas9/guide RNA complex. Application of these programmable regulators is now gaining traction for the modulation of disease-causing genes or activation of therapeutic genes, *in vivo*. Adeno-Associated Virus (AAV) is an optimal delivery vehicle for *in vivo* delivery of such regulators to adult somatic tissue, due to the efficacy of viral delivery with minimal concerns about immunogenicity or integration. However, present Cas9 activator systems are notably beyond the packaging capacity of a single AAV delivery vector capsid. Here, we engineer a compact CRISPR-Cas9 activator for convenient AAV-mediated delivery. We validate efficacy of the CRISPR-Cas9 transcriptional activation using AAV delivery in several cell lines.

## Introduction

As a highly programmable genome editing platform, the CRISPR/Cas9 system has been hailed as a groundbreaking technology to facilitate the treatment of genetic diseases, via modification and correction of detrimental genetic mutations. However, genome editing via CRISPR/Cas9 remains difficult to implement in therapeutic settings, due to concerns surrounding toxicity resulting from double strand breaks, as well as potential off-target mutations.^1–3^ Notable attempts to enhance specificity have been made, including structure-guided engineering of *Streptococcus pyogenes* Cas9 (SpCas9).^4–10^ While enhanced specificity variants eliminate the majority of off target editing, concerns remain as to editing at sites that contain a single mismatch or repetitive sequence.^10^ Additionally, such variants have been shown to trade activity for specificity; room for improvement seemingly remains.^11,12^ Due to these concerns surrounding toxicity and specificity, *in vivo* therapeutic applications based on genetic sequence editing remain tenuous.

Meanwhile, CRISPR/Cas9 activator mediated transcriptional regulation has emerged as a promising alternative strategy, allowing for regulation of gene expression *in vivo*, mollifying concerns of off-target mutations.^13^ Transcriptional regulators have been engineered via direct fusion or recruitment of synergistic, transcriptional activation domains to a nuclease-null Cas9 (dCas9)/gRNA complex. Examples of strong actuators include tripartite dCas9-VP65-p65-RTA direct fusion (dCas9-VPR), synergistic activation mediator (SAM), and repeating peptide array based activator scaffold (dCas9-SunTag).^14–16^

Such activators are now drawing attention, for their potential to treat a variety of diseases where increased expression of a functionally complementary gene can compensate for insufficient activity of a mutated or otherwise disrupted gene.^17^ This has been successfully demonstrated *in vivo* through the delivery of a modified version of the SAM system, in a dual vector Adeno-Associated Virus (AAV) system, for the treatment of a Duchenne Muscular Dystrophy phenotype in mice via activation of *Utrophin* to compensate for mutated X-linked *Dystrophin*.^13^

However, a primary factor in ensuring such strategies will be translatable to the clinic for therapeutic use will be optimization of delivery. Dual vector systems will be more expensive to produce, difficult to titrate stoichiometrically, and cumbersome to work with than a single vector system. Towards the end of pushing transcriptional activators closer to clinical relevancy, we present a rational method of engineering an AAV-compatible Cas9 activator deliverable in a single AAV vector. We systematically modify the design of the dSpCas9-VPR system by employing a small Cas9 ortholog from *Staphylococcus aureus*, identifying complementary truncated functional units of individual activator domains, and validating the use of a compact promoter and terminator. We validate the efficacy of packaging this novel, compact, Cas9 activator into a single AAV vector via delivery to several cell lines, demonstrating robust target gene activation.

## Results

### Engineering a small ortholog activator

While SpCas9 is most commonly employed for genome editing and transcriptional regulation applications, the *Streptococcus thermophiles* Cas9 (St1Cas9) and *Staphylococcus aureus* Cas9 (SaCas9) orthologs are more suitable to packaging in AAV due to their minimal size.^18,19^ While SpCas9 is 4.2 kb long, barely able to fit in an AAV vector, the St1Cas9 ortholog is 3.4 kb while the SaCas9 is 3.2 kb; both can comfortably fit within the 4.7 kb size limit of AAV.

CRISPR/Cas9 activator systems are even more difficult to package within the size limit of an AAV vector. We have previously developed a highly efficient activator, dSpCas9-VPR; however, dSpCas9-VPR is far beyond the packaging capacity of a single AAV vector (standing at 5.9kb).^14^ Other systems including the SAM and SunTag systems suffer from similar issues, and stand well beyond the packaging limit (5.5kb and 7kb respectively.)^15,16^ The primary issue is the Cas9 ortholog used in all of these systems is the SpCas9, which stands at 4.2kb, making it nearly impossible to package itself. As such, if a smaller ortholog could be used in place of the larger Cas9 ortholog in activator systems, this would greatly increase the flexibility of the tool in terms of viral packaging.^19^

Previously, we have validated the St1Cas9 ortholog for its capacity to serve as a modular DNA-binding domain to scaffold the VPR activator on.^14^ However, the PAM of the ST1 ortholog (NNAGAAW) occurs at a relatively low frequency (roughly once every 512 bp) which makes it unattractive for engineering and downstream applications.^18^ To overcome this barrier to a small Cas9 ortholog for *in vivo* delivery with a flexible PAM, the SaCas9 ortholog has been identified as a viable alternative. Previous *in vivo* editing success, validate its ability to edit genes in mouse liver via AAV delivery, and with a more flexible PAM (NNGRRT) which occurs at a higher frequency (once every 64 bp).^19^ As such, we postulated the modular VPR activator could be ported onto the SaCas9 ortholog, similar to St1Cas9, to create an additional orthogonal activator with compact size and flexible targeting properties. We fused the VPR activator to dSaCas9 and targeted it to a TdTomato fluorescent reporter vector, to quantify the relative activity of the novel activator via flow cytometry in comparison with dSpCas9-VPR and dSt1Cas9-VPR. Similar to the dSt1Cas9 activator, we saw retention of the large majority of activity, with only a modest 1.8-fold hit in activity (Figure 1). At the time of completion of this work, this was the highest activation that has been achieved for any activator of such small size, including truncated versions of dSpCas9 which have been known to have significantly reduced activity.^20^ As such, we chose the dSaCas9-VPR system for further engineering.

**Figure 1:**
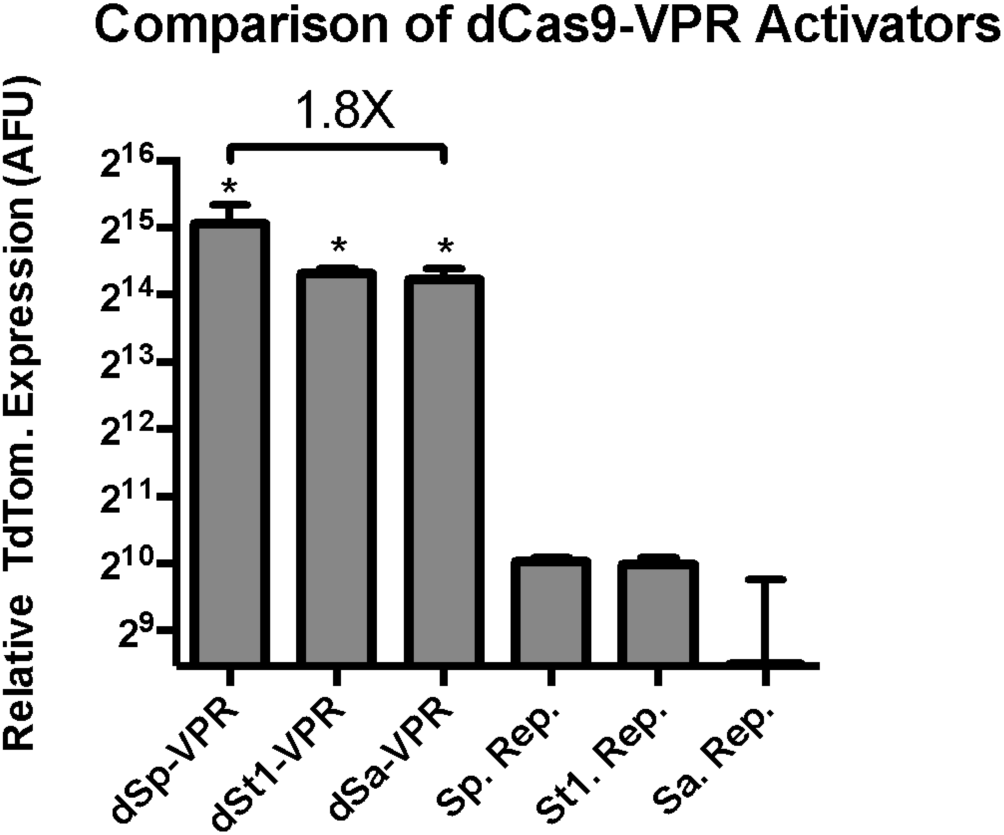
Comparison of VPR activity when fused to small Cas9 orthologs. Transcriptional activation via dCas9-VPR orthologs was performed by fusing a VPR activator to the C-terminus of three nuclease-null dCas9 proteins, *Streptococcus pyogenes (Sp), Streptococcus thermophiles (St1)*, and *Staphylococcus aureus (Sa)*. Cas9 activator orthologs were compared in a fluorescent reporter assay in HEK293T cells. Replacement of dSpCas9 with dSaCas9 to create dSaCas9-VPR leads to a modest 1.8X hit in activity, with a gain of 1kb of genomic sequence space. Error bars indicate median fluorescence ± standard deviation. n=2 biological replicates. (*denotes significance of P =< 0.05 of dCas9-VPR ortholog activation over respective reporter only control via one-tailed t-test.)

### Identification of minimal activator domains and assembly of a robust compact activator

Even with the 1kb gain in sequence space from replacing dSpCas9 with dSaCas9, the entire dSaCas9-VPR tool stands at 4.9 kb to express the dCas9/activator effector alone. With the packaging limit of AAV being 4.7 kb, this tool is impossible to package, let alone with the necessary constitutive promoter, poly-adenylation signal, and sgRNA expression cassette. The most common constitutive promoter for *in vivo* expression is the 600 bp CMV promoter, the poly- adenylation signal used to express SaCas9 for *in vivo* editing is the bGHR signal of 250 bp, and the standard sgRNA expression cassette consists of the 250 bp RNA pol II U6 promoter and 100 bp sgRNA sequence. As such, additional minimization of composing elements is required.

We hypothesized that the activation domains (p65 and RTA, originally identified to build the VPR activator) could be truncated to isolate only necessary elements for activator domain function. We performed rational truncations on p65 and RTA to identify several minimal domains that retained activity. Furthermore, we hypothesized that the truncated domains might retain their ability to induce high levels of gene expression, comparable to the full length VPR activator, when fused together as a synergistic unit.

The p65 activator contains two well-known transcriptional activation domains (TADs) at 415-459aa and 536-533aa. Similarly, RTA contains a known TAD from 530-691aa. However, truncations of these activation factors have not previously been attempted. As such, we performed serial truncations of p65 and RTA from either the N or C-terminus, to validate whether a smaller activator could be recovered (Figure 2). We then fused each of these activators to dSpCas9, and screened for relative activity in our previously described TdTomato reporter system. We identified several smaller truncated activators, which retained a large percentage of activity, including the two smallest activators p65 (100aa-261aa) and RTA (125-190aa). The two truncated effectors retained the majority of their full- length activity, respectively, while recovering 675 bp in sequence space (Figure 3). In order to validate that a combination of truncated factors still performed synergistically to activate targeted genes effectively, we combined the most promising truncated domains into novel truncated versions of the original VPR activator, and fused these minimized activator combinations to the SaCas9 ortholog. We delivered the novel compact activators to N2A cells, targeting a panel of genes including *Actc1, Acta1*, and *Hbb*. The two smallest truncated activators retained essentially all activity when combined in the complete tool and compared to full length SaCas9-VPR, while bringing the size of the activator tool itself down from 5kb to 4.3kb, well within the packaging limit of AAV (Figure 4).

**Figure 2:**
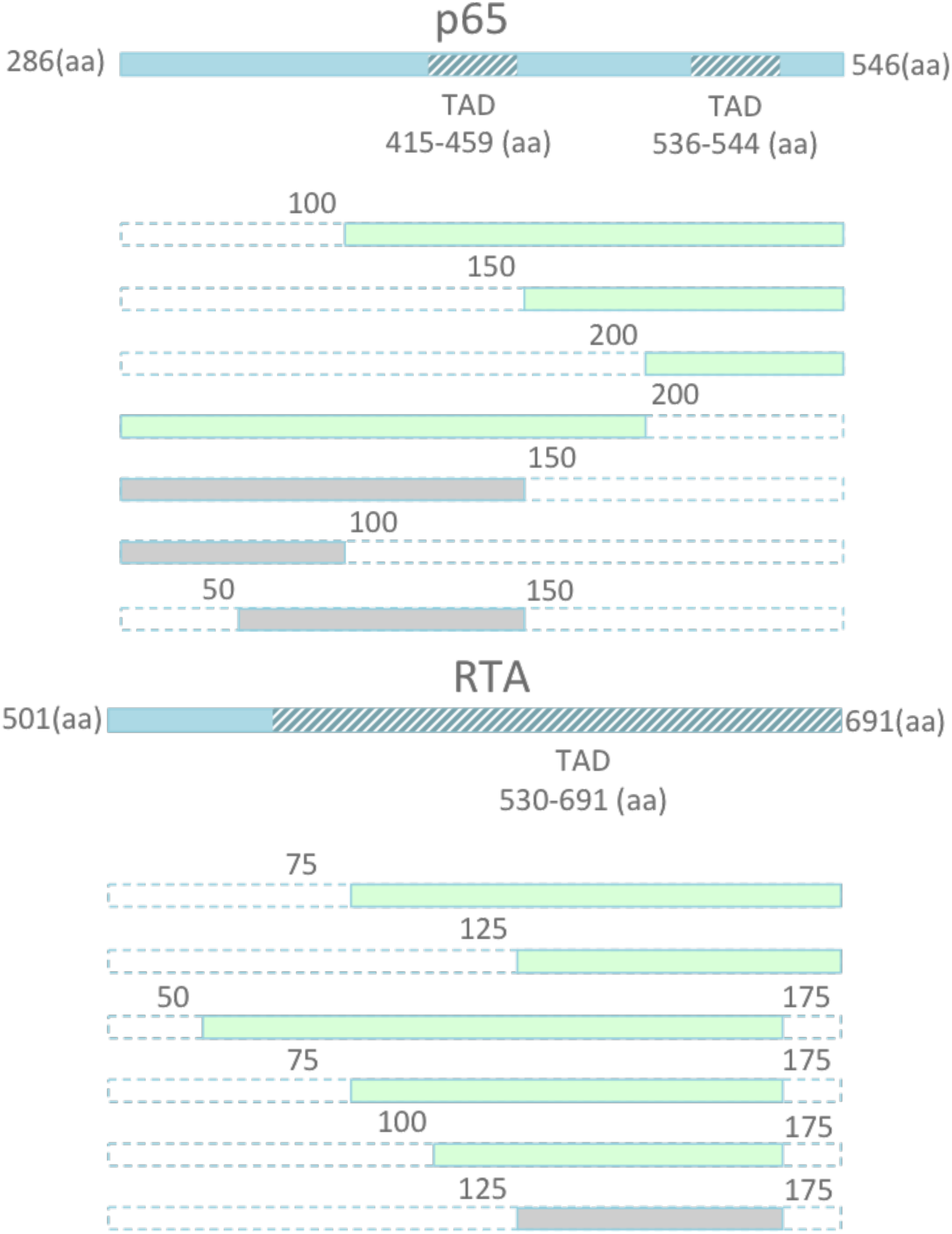
Schematic illustration of TADs located within p65 and RTA. A) Schematic of serial truncations of the full length 261 amino acid (aa) p65 from the N terminus, C terminus, and both are fused to the C-terminus of dSpCas9. The retained aa positions of the new domain are indicated by the numbers surrounding each domain depicted in green. For example, p65 (100-261) indicates a truncated p65 domain with the first 99 aa removed, and the remaining domain retained.

**Figure 3:**
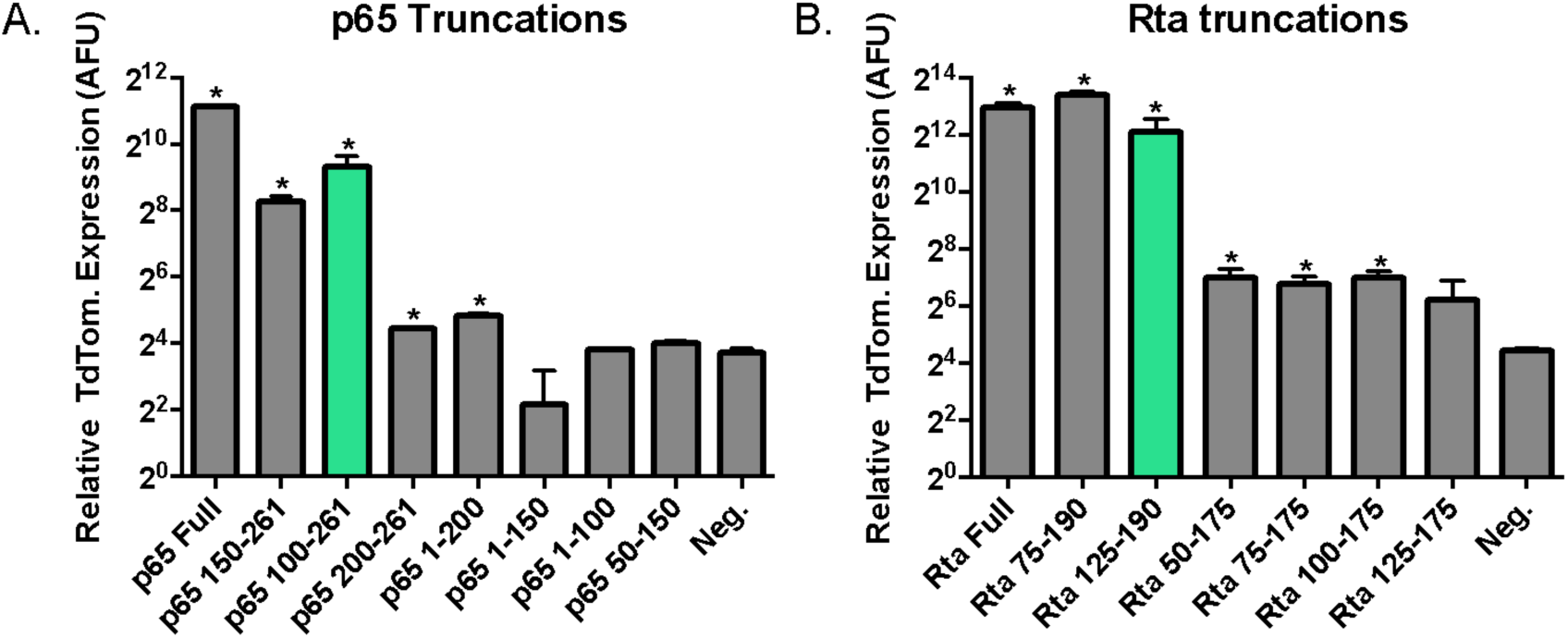
Screen for truncated p65 and RTA domains that retain activity. A) Serial truncations of the full length 261 amino acid (aa) p65 from the N terminus, C terminus, and both are fused to the C-terminus of dSpCas9. The retained aa positions of the new domain are indicated by range of numbers following the domain name. For example, p65 (100-261) indicates a truncated p65 domain with the first 99 aa removed, and the remaining domain retained. Each activator, containing full length p65 or each of the 7 truncated versions of the domain, are compared in a fluorescent reporter assay in HEK293T cells. Truncation of p65 to produce p65 (100-261) lowers activity a modest 3 fold, while allowing for a gain of 300bp of sequence space. B) Similarly, removal of the first 125 aa of the RTA domain lead to a negligible 2 fold reduction in activity, while allowing for a gain of 375bp of sequence space. Additional truncations retained activity while allowing for a reduction in length, including p65 (150-261) and RTA (75-190). Error bars indicate median fluorescence ± standard deviation. n=2 biological replicates. (*denotes significance of P =< 0.05 of activator over reporter only control via one-tailed t-test.)

**Figure 4:**
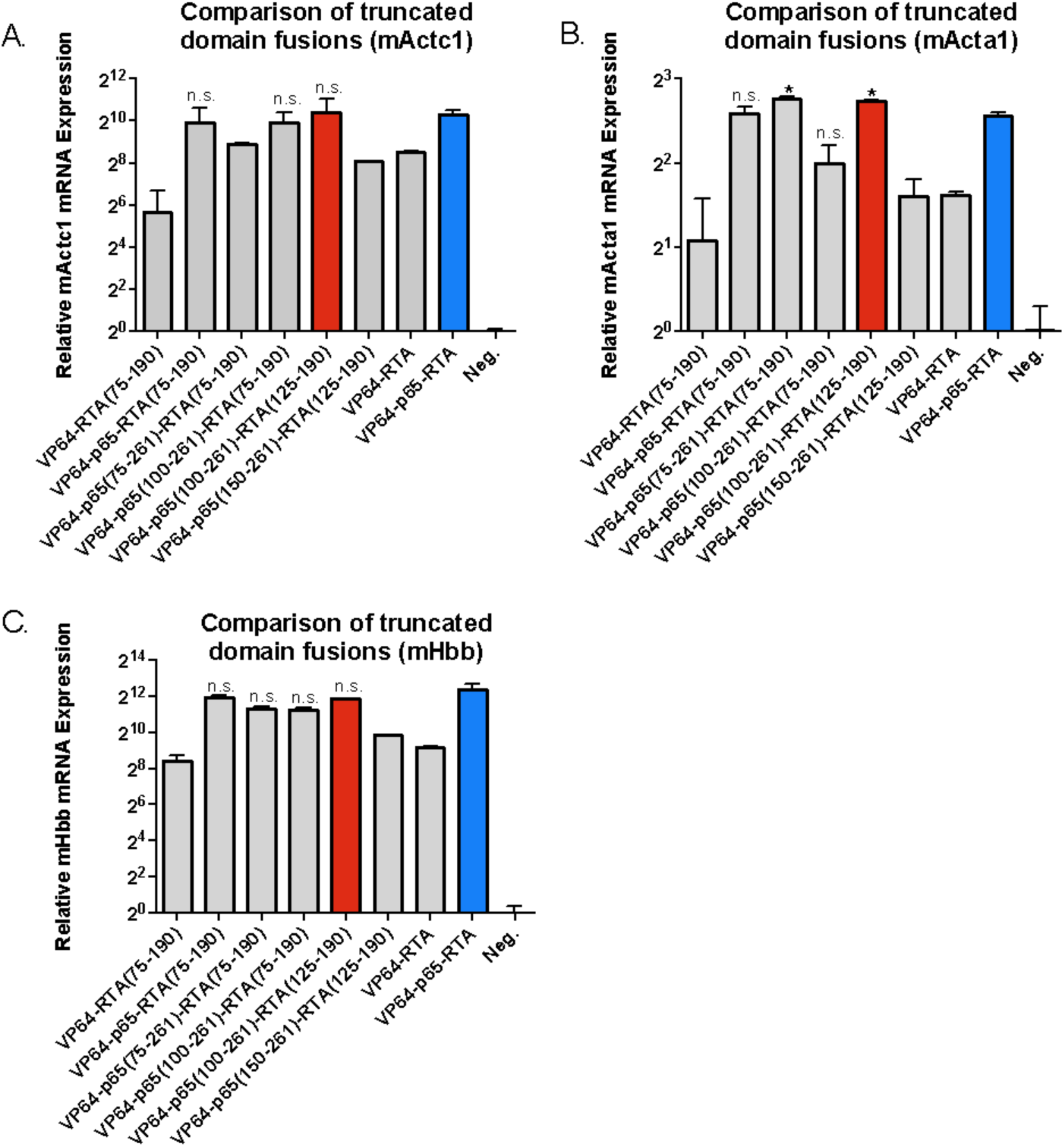
Comparison of combinations of truncated activation domains fused to C terminus of nuclease-null SpCas9.A) Both p65 and RTA in the synergistic VP64-p65- RTA activator are replaced with their respective truncated versions. The new set of smaller activators are targeted to the endogenous mouse *Actc1* gene in Neuro-2A cells. The VP64-p65-RTA(75-190), VP64-p65(100-261)-RTA (75-190), and VP64- p65(100-261)-RTA(125-190) show no significant loss in potency compared to the full length VP64-p65-RTA activator. The smallest activator that retains potency, VP64-p65(100-261)-RTA(125-190), affords a 675 bp gain in sequence space. B) The new set of activators are targeted to the endogenous mouse *Acta1* gene in Neuro-2A cells. Both VP64-p65(75-261)-RTA(75-190) and VP64-p65(100-261)-RTA(125- 190) show a surprising and significant increase in potency over the full length VP65- p65-RTA activator. C) The new set of activators are targeted to the endogenous mouse *Hbb* gene in Neuro-2A cells. Four of the seven novel activators show no significant loss in activity relative to the full length activator, including the smallest new tool, VP64-p65(100-261)-RTA(125-190). Error bars indicate mean expression ± standard deviation. n=2 biological replicates. (*denotes significance of P =< 0.05 of activator when compared to full length activator via one-tailed t-test, n.s. denotes no significance.)

### Validation of novel mini activator against gold-standard

We were able to reduce the size of the Cas9 activator from the original dSpCas9-VPR tool, standing at 5.9 kb, down to a miniature yet potent dSaCas9-VPR employing the p65 (100-261aa) truncation and the RTA (125-190aa) truncation, leading to an effective and compact tool of 4.2kb – well below the packaging limit of AAV. While the dSa-VPR miniature activator is below the packaging limit of AAV, one other activator had previously been identified in the literature that is below the packaging limit of AAV, and that is the dSa-VP64 activator component of the multifunctional SAM tool.^15^ To asses the relative efficacy of the dSa-VPR miniature activator over the dSa-VP64 tool, we targeted both tools to a set of 4 genes and compared relative levels of induction, and demonstrated a 24 to 2100-fold increase in activity on four genes in N2A cells, *Cdr1as, Neurog2, Actc1, and Hbb* (Figure 5). Confirming dSa-VPR mini. as the smallest effective Cas9 activator available, we proceeded to identify suitable expression elements for packaging and delivery via AAV.

**Figure 5:**
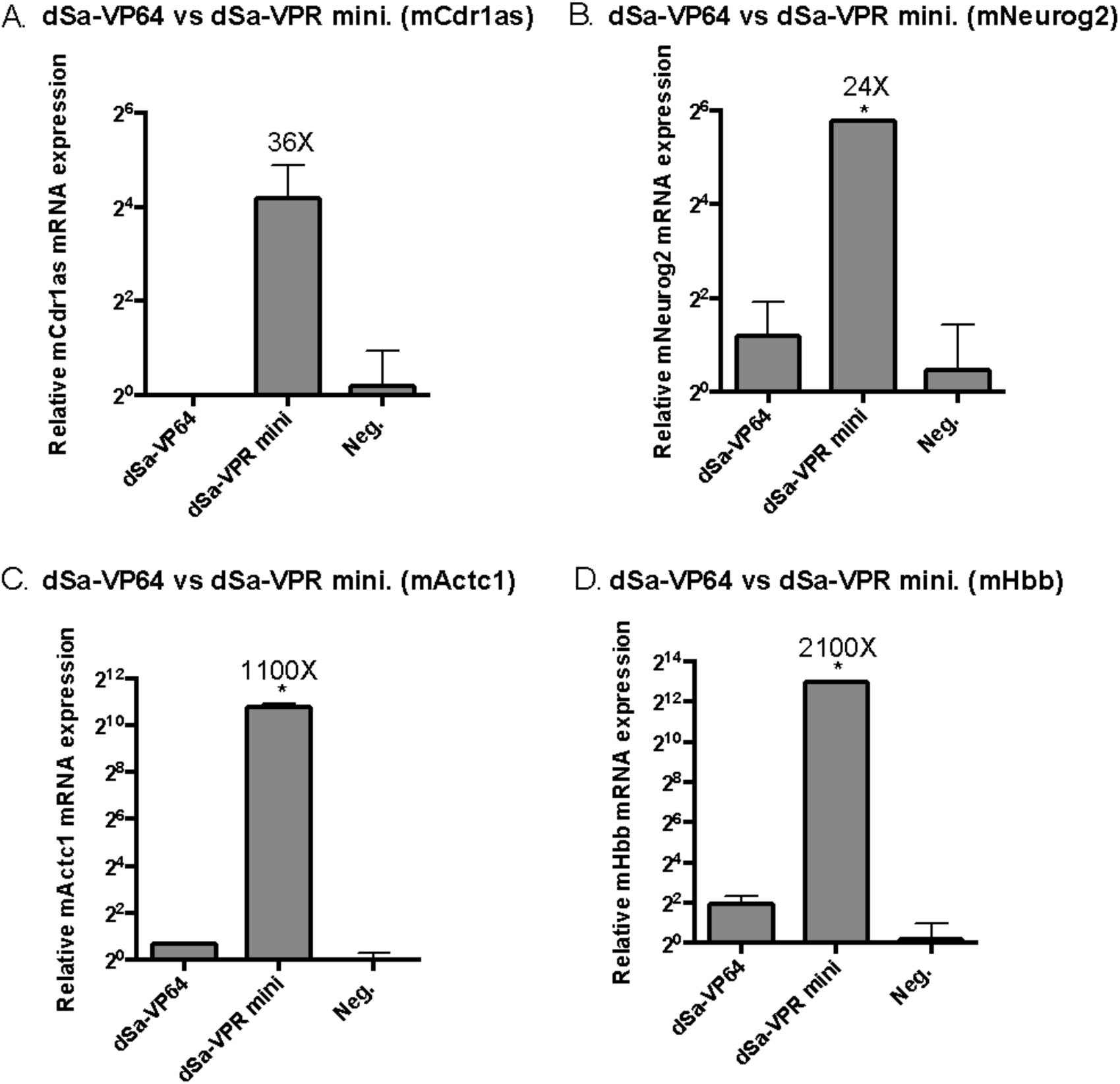
Gene activation with dSa-VPR miniature vs dSa-VP64. A) Both dSa-VP64 and the dSa-VPR miniature activator are targeted to the mouse *Cdr1as* lncRNA gene in Neuro-2A cells. The dSa-VPR miniature activator performs 36-fold better than the other small activator dSa-VP64, however the median is not significantly different from the median level of activation with dSa-VP64, due to a large variance. B) Both tools are targeted to the mouse gene *Neurog2* and dSa-VPR miniature performs 24X better than dSa-VP64. C) Both tools are targeted to the mouse gene *Actc1* and dSa- VPR miniature performs 1100X better than dSa-VP64. D) Both tools are targeted to the mouse gene *Hbb* and dSa-VPR miniature performs 2100X better than dSa-VP64. Error bars indicate mean expression ± standard deviation. n=2 biological replicates. (*denotes significance of P =< 0.05 of dSa-VPR miniature over dSa-VP64 via one- tailed t-test.)

### Identification of effective compact promoters

While the activator tool is below the packaging limit of AAV, standing at 4.2kb, expression elements are necessary for driving transcription and termination of the tool to be expressed. The final design requires the incorporation of a 350 bp guide RNA expression cassette, leaving roughly 150 bp for promoter and termination elements. As such, we screened several short promoters, and identified two small constitutive promoters that would allow for high levels of dSa-VPR miniature expression, without compromising the tool’s ability to activate genes as compared to when expressed off of the gold-standard CMV promoter. The two promoters, SCP1 and EFS, are 80 and 200 bp long respectively, compared to CMV which is 600 bp in length.^21,22^ SCP1-dSa-VPR mini., EFS-dSaVPR-mini., CMV-dSa-VPR mini., and CMV-dSa-VP64 were targeted to both *Neurog2* and *Actc1* in N2A cells and compared for relative expression of the SaCas9 gene as well as the targeted gene. The SCP1 and EFS based designs retained the large majority of expression; enabling equivalent expression of the targeted genes relative to the CMV based design (Figure 6). With the use of the SCP1 promoter, a remainder of 70 bp of space was available for a terminator sequence.

**Figure 6:**
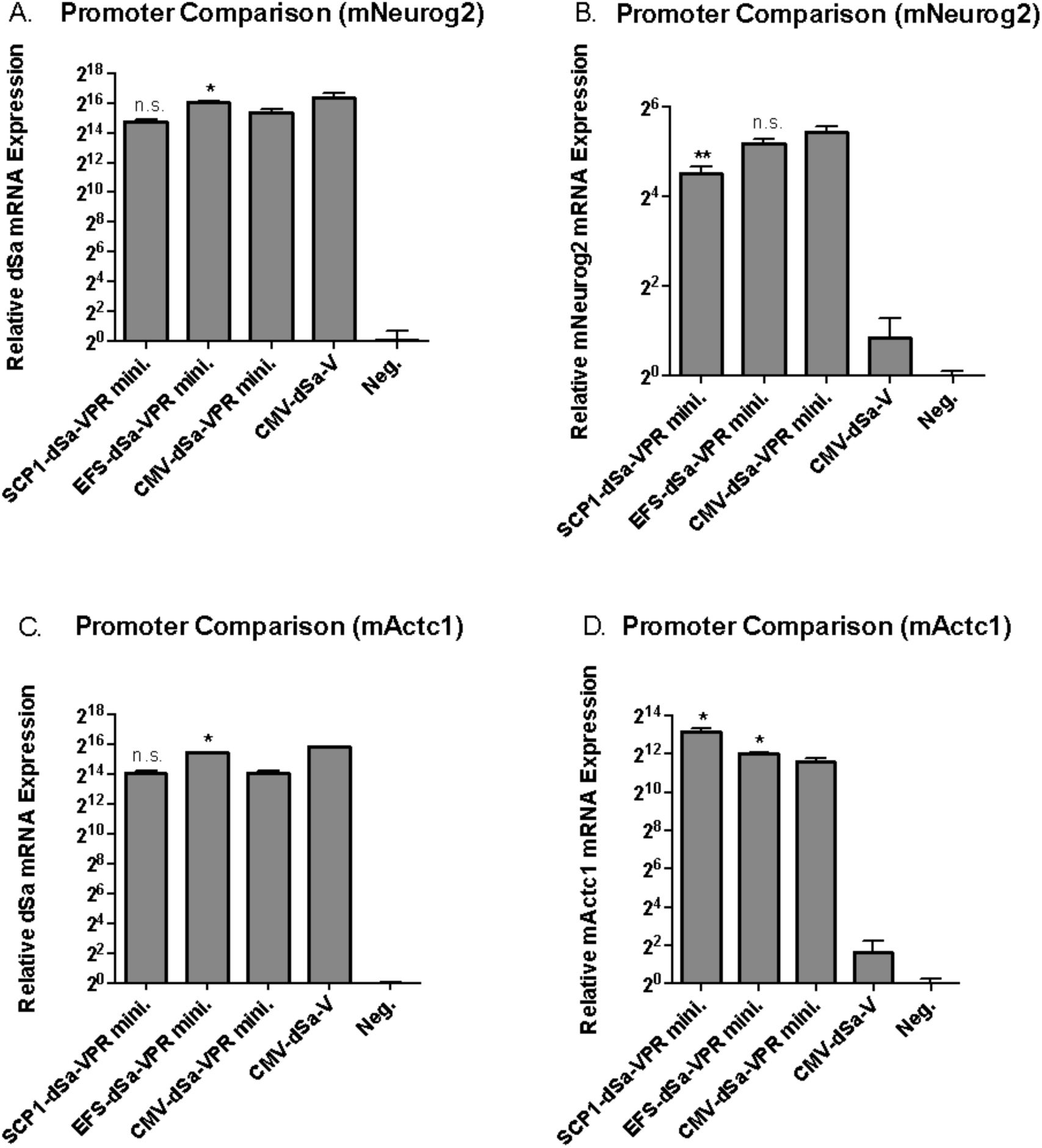
Comparison of novel SCP1 and EFS promoters. A) The dSa-VPR miniature activator is expressed off of a SCP1 (Super Core Promoter 1) promoter, EFS (Elongation Factor Short) promoter, or a CMV (Cytomegalovirus) promoter and targeted to the mouse gene *Neurog2* in Neuro-2A cells. The targeted cells are assayed via qPCR for expression of the dSaCas9. While SCP1 allows for an expression on level with CMV, the EFS promoter significantly enhances the amount of activator transcript present, compared to CMV. B) The same population of cells in A) are assayed for expression of the *Neurog2* target gene, use of the SCP1 promoter reduces efficiency roughly two-fold relative to the CMV promoter, while the EFS promoter essentially maintains efficiency relative to CMV. C) The set of three dSa- VPR miniature expression cassettes, driven by SCP1, EFS, or CMV, are targeted to the mouse gene *Actc1*. The cells are assayed for expression of the dSaCas9 via qPCR. No significant loss in expression is observed due to the use of SCP1 relative to CMV, and again, an enhancement in expression is observed due to the use of EFS to drive expression as opposed to CMV. D) The same set of cells in panel C) are assayed for expression of the target gene *mNeurog2*. Both the SCP1 and EFS driven activators show an enhancement in efficiency relative to the CMV driven cassette. Error bars indicate mean expression ± standard deviation. n=2 biological replicates. (*denotes significance of P =< 0.05 of a short promoter driven activator relative to the CMV driven cassette via one-tailed t-test. **denotes significance of P=<0.05, in the opposite direction, with the short promoter driven activator performing at a lower efficiency than the CMV driven cassette.)

### Identification of effective minimal terminators

Similarly, to identify previously under-utilized termination compact termination sequences, we screened several poly adenylation signals, and identified an extremely short dual function polyadenylation (pA) signal, containing a stop codon signal, that is efficient for termination of gene transcription and translation.^23,24^ The short 17 nt terminator was previously shown to work efficiently *in vivo*, and a dual repeat (34 nt) version was shown to work comparatively efficiently to several other polyadenylation signals including the popular SV40 polyA sequence.^23^ We compared the a single repeat sNRP-1 polyA signal, dual repeat version, the synthetic polyA signal, and the commonly used bGHR polyA signal for ability to stabilize activator transcripts and therefore enable downstream activation of targeted genes.^25^ We targeted the designs to several target genes including *Actc1, Hbb, and Neurog2* in N2A cells and assayed for both dSa and target gene expression. Given size constraints and ability to retain activation of the targeted genes, the 2X- sNRP-1 terminator was chosen (Figure 7).

**Figure 7:**
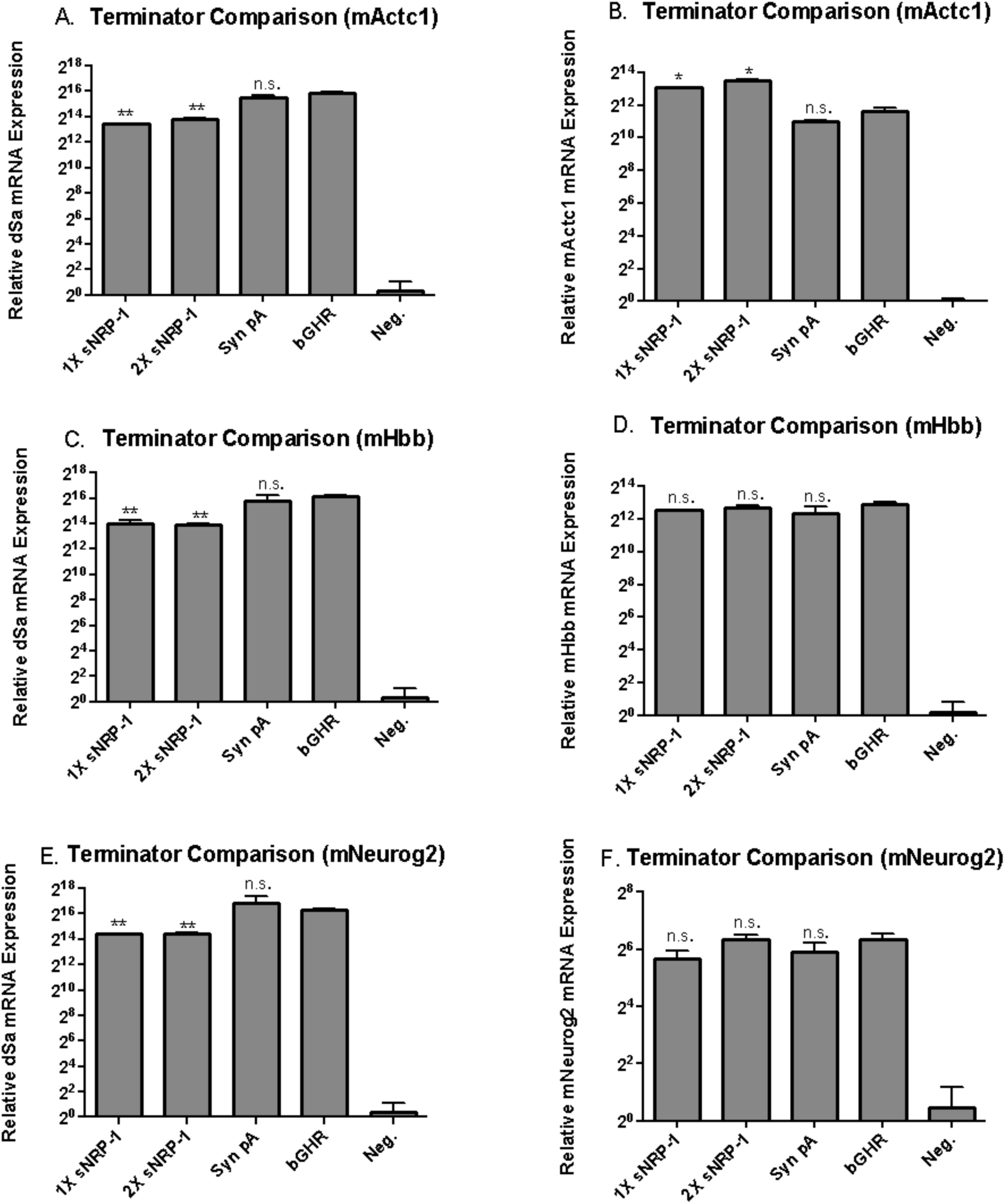
Comparison of extremely short poly adenylation sequences. Short 17nt sNRP-1, a 34nt dual sNRP-1, a 50nt synthetic, and a 250 nt bGHR terminators were cloned to the end of the dSa-VPR miniature expression cassette. A) Each construct was targeted to the mouse *Actc1* gene via transfection in to Neuro-2A cells, and compared to the commonly used bGHR signal via qPCR for presence of dSaCas9 transcript, 48 hours post transfection. Both the extremely short 1X sNRP-1 and 2X sNRP-1 terminators were able to stabilize transcript expression, albeit at a 5-fold reduction in transcript number relative to the bGHR signal. B) Despite mediating a reduction in transcript number, the 1X sNRP-1 and 2X sNRP-1 poly adenylation signals enable a surprisingly 3-fold increase in target *Actc1* expression, indicating that the lower transcript number does not necessarily reduce the efficiency of dSa- VPR miniature mediated activation of the target. C) Similar to panel A, each construct is targeted to the mouse *Hbb* gene and compared for expression of the dSaCas9 transcript. Both the 1X sNRP-1 and 2X sNRP-1 terminators show a significant reduction in transcript number relative to the bGHR terminator. D) However, when the cells are assayed for target *Hbb* expression, there is no significant difference in target expression between the 3 short terminators and the bGHR signal. E) The constructs are targeted to the mouse gene *Neurog2* and compared for dSaCas9 transcript expression. Again, a reduction in transcript number is observed with the 1X sNRP-1 and 2X sNRP-1 terminators, relative to the bGHR signal. F) Yet, when the cells are assayed for *Neurog2* target expression, no significant difference is found in gene expression level between the 3 terminators compared to the bGHR signal. Error bars indicate mean expression ± standard deviation. n=2 biological replicates. (*denotes significance of P =< 0.05 of a short terminator mediated expression relative to bGHR via two-tailed t-test. **denotes significance of P=<0.05, in the opposite direction, with the short terminator stabilized activator expressing lower level than the bGHR stabilized tool.)

### Activation of endogenous genes with single vector system

With a small 3.2kb *Staphylococcus aureus* Cas9, a 1kb miniature yet potent VPR activator, constitutive yet unprecedentedly small 80 bp SCP1 promoter, and a novel extremely short dual function transcription and translation 34 bp 2X sNRP1 terminator, we reached a theoretical activator expression cassette totaling just above 4.3kb – well below the packaging limit of AAV. In order to create a single vector system, we combined all elements, alongside a 350 bp sgRNA expression cassette, to create a 4.75 kb tool ready for packaging in to AAV (Figure 8).

**Figure 8:**
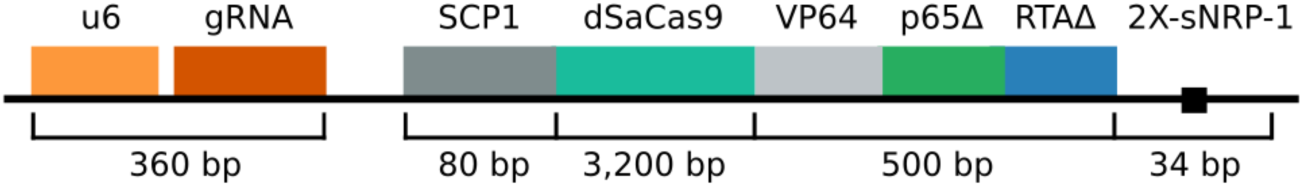
Diagram of full single vector activator system, including sgRNA expression cassette, SCP1 promoter, dSaCas9-VPR miniature activator, and 2X-snRP-1 within a single 4.75kb construct.

We validated the new single vector cassette for the ability of all elements to efficiently work together by comparing the design to the only other activator that will currently fit within the packaging limit of AAV – the dSa-VP64 tool. Due to the smaller size of the dSa-VP64 tool, we were able to include full-length CMV promoter and bGHR signals, along with the sgRNA expression cassette, making the most of the space available when using the smaller activator. We then compared the two single vector tools for ability to activate several target genes via transfection, to validate the enhanced potency of our novel single vector activator expression system, relative to the only other small activator available using gold standard expression elements on three target genes *Actc1, Neurog2,* and *Hbb* in N2A cells, and show that the novel compact tool activates target genes 104, 110, and 680 fold above background, corresponding to 104, 18, and 680- fold better than the only other single vector tool available (Figure 9). However, while the tool had been extensively validated in the N2A neuron derived cell line, eventual downstream applications would require activity in a variety of cell lines and tissue types. As such, we validated the tool in three additional diverse cell lines including mouse Hepa1-6 hepatocarcinoma, GC-1 spermatogonial, and C2C12 muscle myocyte lines (Figure 10).

**Figure 9:**
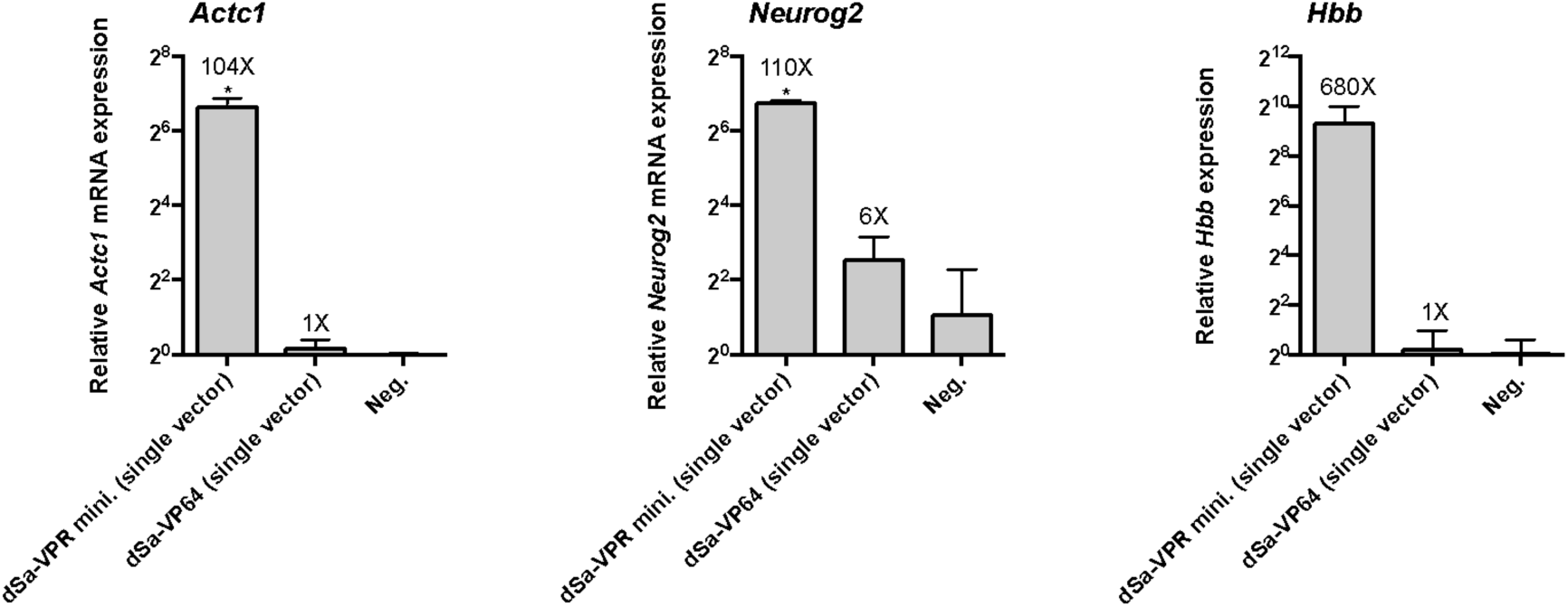
Comparison of single-vector activator designs. A) The new activator construct and the gold standard vector are targeted to the mouse *Actc1* gene. Vectors are transfected in to Neuro-2A cells, and assayed for target expression after 48hours. The new design performs 104-fold better than the standard. B) The two tools are targeted to the mouse *Neurog2* gene, and the new design performs 18-fold better than the standard. C) Both activators are targeted to the mouse *Hbb* gene, and the new design performs 680-fold better than the standard. Error bars indicate mean expression ± standard deviation. n=2 biological replicates. (*denotes significance of P =< 0.05 of improvement in target activation with the new design relative to the standard, via one-tailed t- test.)

**Figure 10:**
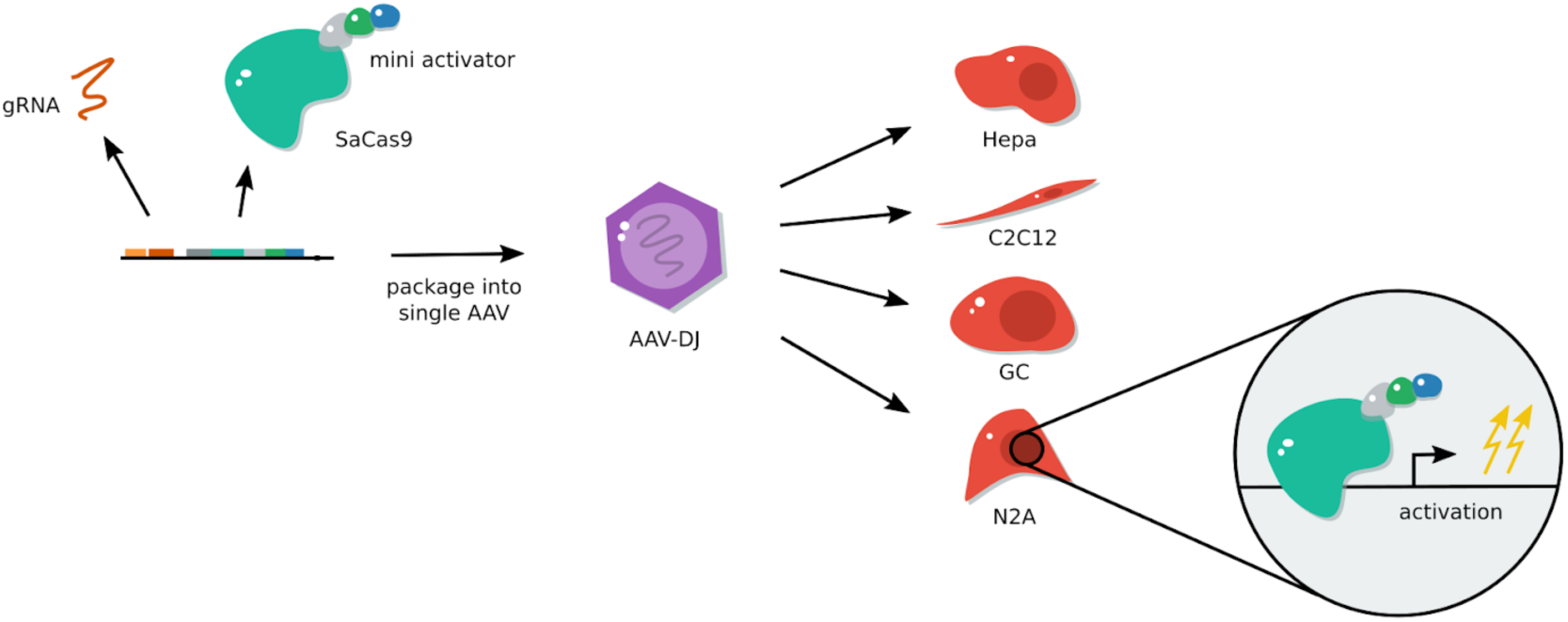
Schematic of gRNA, and dSaCas9-VPR miniature activator, packaged into a single AAV and delivered to several cell lines, including Hepa 1-6, C2C12, GC-1 and N2A lines.

### Validation of single vector activator in several additional mouse cell lines

The new tool performs far better than the alternative standard activator tool that would be available for packaging in to an AAV expression cassette. While many applications of *in vivo* transcriptional regulation will undoubtedly occur in neuronal cell types, we wished to validate the efficacy of the SCP1 promoter, the novel miniature Cas9 activator, the compact sNRP-1 terminator, and guide RNA expression system in a variety of cell lines, to confirm the relevance of the tool for use in other tissue types. Thus, we targeted our novel activator to a panel of genes in 3 additional cell lines, C2C12 mouse myocytes, GC-1 mouse spermatogonial cells, and Hepa 1-6 mouse hepatocarcinoma cells, as well as Neuro-2A cells. Since the target genes will be present in a variety of epigenetic states when targeted in different cell types, and it is possible that genomic rearrangements in immortalized cell lines might even lead to a loss of the targeted locus in a new cell line, we did not expect all targets to activate in every cell line. However, at least one target was activated extremely well in each cell line, validating that each element is working in all four lines, albeit at different efficiencies depending on the locus (Figure 11).

**Figure 11:**
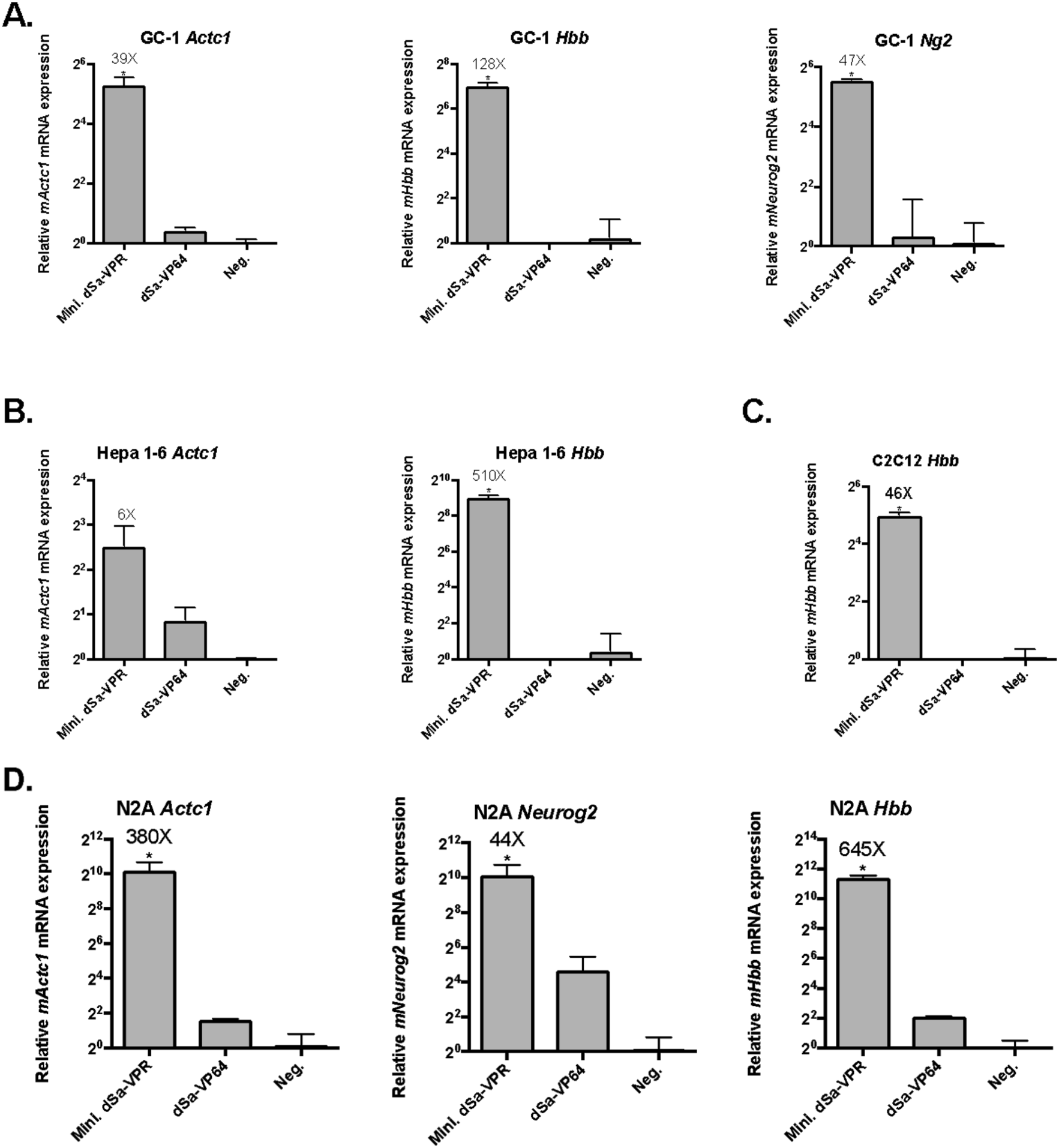
Validation of new activator tool in 4 cell lines. A) The novel tool is targeted to the three mouse genes in GC-1 spermatogonial cells. The *Actc1, Hbb,* and *Neurog2* targets are highly induced when activated by dSaCas9-VPR mini. relative to the gold standard small activator. B) Two genes are successfully targeted in mouse Hepa 1-6 hepatocarcinoma cells; *Actc1* is modestly up-regulated, *Hbb* is strongly induced. C) The target *Hbb* is highly induced in C2C12s. D) The three genes are targeted in mouse Neuro-2A cells and are all highly up-regulated, as previously observed. (*denotes significance of P =< 0.05 of improvement in target activation with the new Mini. dSa-VPR design relative to the dSa-VP64 standard, via one-tailed t-test.)

### Packaging single vector activator in AAV

With the tool validated in 4 separate mouse cell lines *in vitro*, we packaged the new Cas9 activator expression system in to high titer AAV. AAV has 12 serotypes available for packaging, with a variety of tropisms specific to targeting a variety of tissue types including neuronal cells, muscle, liver, retinal cells, etc. However, an alternative serotype AAV-DJ has been developed, which allows for relatively efficient delivery to *in vitro* cell lines. In order to more rapidly validate that the tool packaged efficiently, could be delivered via AAV, and could mediate activation of target genes when packaged in AAV, we chose to initially package a tool targeting the mouse *Actc1* gene in to AAV-DJ. To validate the packaging efficiency, we titered the vector by running qPCR on the packaged virus alongside qPCR of linearized vector standards, with defined copy numbers. Additionally, we denatured the AAV virus, and ran the packaged genomes on a 1% Agarose E-gel with SYBR Gold stain, which visualizes single stranded DNA, alongside cut vector standards. Via qPCR, we detect a titer of roughly 10^12 total viral particles isolated for the mActc1 targeting activator tool, and see a roughly 5kb clean band on the electrophoresis gel, indicating that the vector has been correctly packaged (Figure 12).

**Figure 12:**
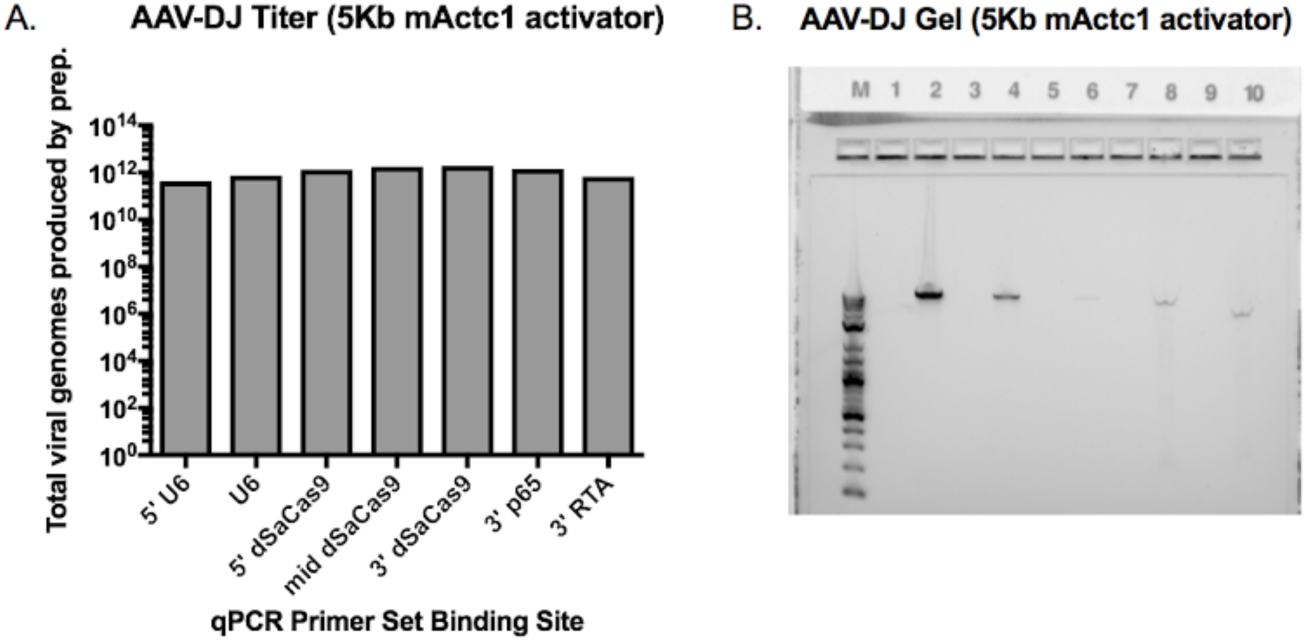
Validation of *Actc1* targeting tool for AAV-DJ packaging. A) qPCR of 7 different portions of the new activator tool, progressing from the 5’ end to the 3’ end of the packaged genome. Viral genomes are calculated via standard curves generated from standard qPCR curves produced from qPCR of cut vector standards, with known copy numbers of the new activator tool expression cassette. B) Electrophoresis agarose gel with SYBR Gold stain for single and double stranded DNA. Lane M is Log-2 ladder, lane 2 is 200 ng of cut vector (7kb), lane 4 is 50 ng of cut vector (7kb), lane 6 is 1ng of cut vector (7kb), lane 8 is 10^9 AAV-DJ viral particles packaged with *Actc1* targeting Cas9 activator tool (5kb, including ITRs), lane 10 is 10^9 AAV-DJ viral particles packaged with CAG-GFP expression cassette (3kb, including ITRs.)

### Delivery of single vector activator via AAV to activate target genes in vitro

To validate the packaged *Hbb* targeting activator tool, the vector was packaged through a modified small-scale preparation protocol in AAV-DJ. The virus was approximately 1e7 vg/uL, and we delivered roughly 1e8 particles to C2C12, Hepa 1-6, GC-1 cells seeded in a 24-well plate (Figure 13). The virus was toxic to Neuro-2A cells, preventing collection of RNA for downstream processing of expression data. After 9 days, the *Hbb* target locus was successfully up-regulated by AAV-DJ mediated delivery of the compact activator in all three lines, ranging from 54 to 149-fold up-regulation, corresponding with a 9 to 300-fold improvement over the only other available activator to be packaged in a single AAV vector. Thus, we successfully validated the efficacy of our single vector dSaCas9 miniature activator, for delivery in a single vector AAV, enabling activation of a target gene in several cell lines.

**Figure 13:**
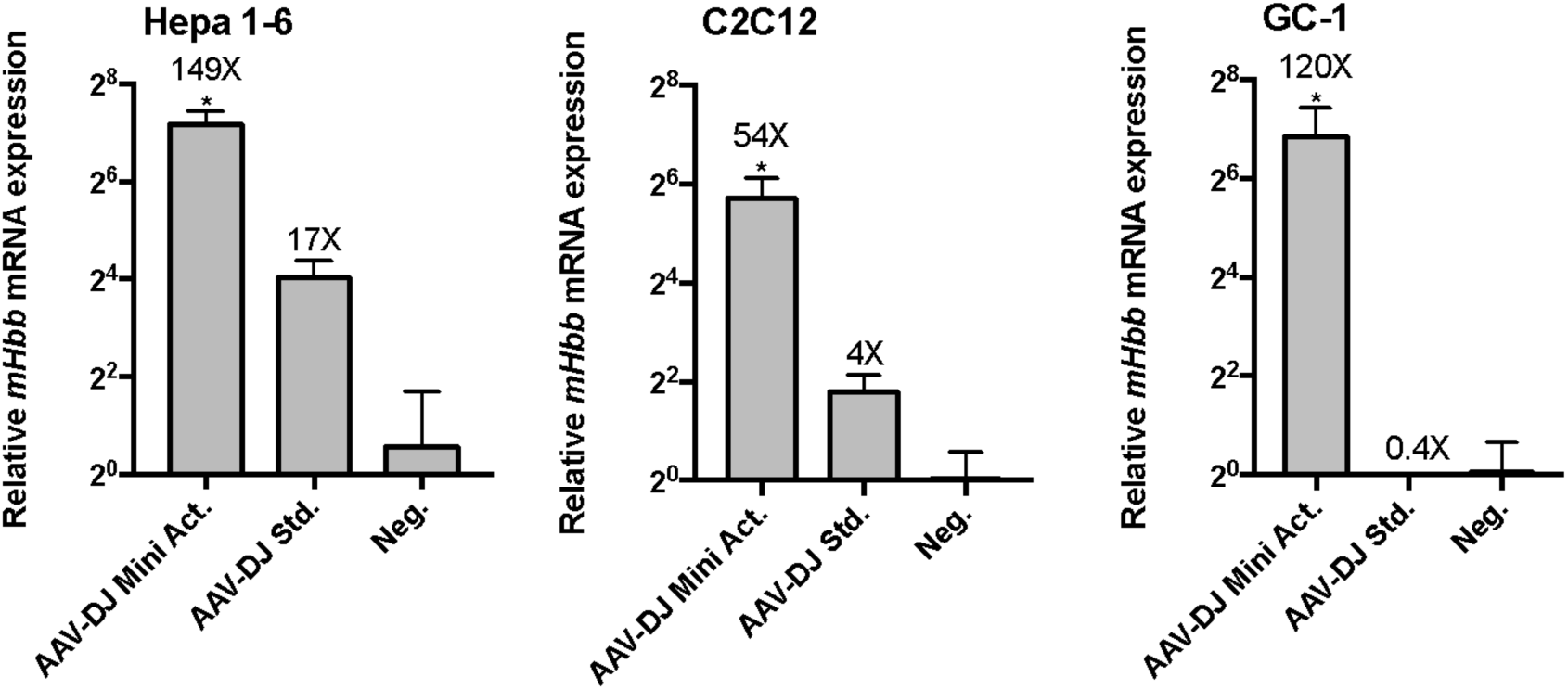
Delivery of AAV-DJ with *Hbb* targeting activator tool to 4 cell lines *in vitro*. A) Roughly 10^9 viral particles of the packaged activator tool in AAV-DJ are delivered to C2C12, Hepa1-6, C2C12, and GC-1 cells, and assayed for *Hbb* target expression after 9 days. As expected from transfection experiments, the *Hbb* target gene is up-regulated in all three cell lines, between 54 and 149 fold above background, corresponding to 9-300 fold above that achieved with the dSa-VP64 based standard. (*denotes significance of P =< 0.05 of improvement in target activation with the new Mini. dSa-VPR design relative to the dSa-VP64 standard, via one-tailed t-test.)

## Discussion

Directed regulation of therapeutic gene targets via AAV mediated delivery CRISPR/Cas9 activators, holds potential both as an exciting method for enabling gain of function studies in *in vivo* disease models, and as a novel clinical method for treatment of several currently intractable genetic disorders. As recently demonstrated with this method, it is now possible to ameliorate phenotypes associated with mouse models of Duchenne Muscular Dystrophy, Diabetes, and Acute Kidney Disorder through activation of compensatory, differentiation inducing, or more generally protective therapeutic genes. ^13^

However there are many other genetic disorders that may benefit, at least in terms of basic research, from such a tool. Disorders related to genomic imprinting, such as Angelman Syndrome and Prader-Willi Syndrome, occur due to mutations in either a paternal or maternal allele specifically; while a functional copy of the mutated gene of interest remains intact on the other “imprinted” or silenced allele. In such cases, it may be plausible to induce expression of the functional, yet epigenetically silenced allele.^26^ Additionally, disorders due to haploinsufficiency, may be addressed by overexpression of a single functioning allele. Expanding upon this paradigm, disorders caused by expanded nucleotide repeats which result in epigenetic silencing of a normally functioning, necessary gene – as is the case in Fragile X Syndrome and Freidrich’s Ataxia – may benefit from attempts to revert the silenced gene to an active state.^27,28^ Finally, several disorders exist, akin to Duchenne Muscular Dystrophy, where mutations in structural proteins cause debilitating and life threatening disorders; however, in select cases, there exist separately regulated fetal variant or separate isoform of the diseased gene, which can offer compensatory functionality if induced. Examples of such disorders include Nemaline Myopathy, Sickle Cell Disease, as well as Spinal Muscular Atrophy. ^29–31^

Many of these highlighted disorders are due to mutations that affect cellular function in difficult to reach, or large areas of tissue, that will be difficult to deliver a virus to. In such cases, where neurological tissue, or expansive, difficult to reach tissue must be studied or treated, a dual vector AAV system may be unnecessarily cumbersome to administer in a broad range of experimental and clinical settings as opposed to a single vector virus.^32^ In such cases, the development of single vector variants of activators serves well in terms of utility for future applications. As such, while this manuscript was under preparation, a similar work was published demonstrating feasibility of delivering a Cas9 activator in a singular AAV vector.^33^

Looking forward, Cas9 activator mediated activation remains an exciting alternative method for the treatment of genetic disorders; yet, an AAV based system alone does not solve all potential related issues of toxicity or delivery. While the AAV system itself has now generally been considered benign and effective for delivery in clinical setting, Cas9 itself has yet to be thoroughly vetted for toxicity clinically.^34^ Early indications have shown immunogenic responses to Cas9 itself, and careful research must be conducted to vet such concerns.^35^ As such, the activator presented here may instead be used with ZFs or TALEs as well, and remain easily within the packaging limit; possibly allowing for more flexibility in terms of expression elements such as promoters, terminators, or fluorescent markers. There are some advantages to these platforms, such as a reduced immunogenicity, size, and previous clinical trial data that make them potentially attractive for human gene regulation therapy applications.^36^ Overall, we remain optimistic about the role of AAV CRISPR/Cas9 based activator systems, particularly in terms of their role in basic research, and intently await further studies demonstrating their appropriateness for use in clinical settings.

## Author Contributions

S.V. and A.C. conceived of the study. S.V., and J.C. designed and performed experiments and interpreted data with support from A.C., R.X., N.J.V., and L.Q. W.T.P., L.H.V., A.C., and G.C. supervised the study. S.V. wrote the manuscript with support from J.C. and all other authors.

## Acknowledgements

We thank the Gene Transfer Vector Core for producing a subset of the vectors used in this work. Additionally, we thank the Zhang lab for their gift of an AAV vector containing nuclease null SaCas9. Additionally, we thank gracious former and current members of the Church lab, Kyle Cromer, Wei Leong, and Noah Davidsohn for their support in large scale AAV production.

## Materials and Methods

### Vectors used and designed

Combined activation domains were cloned using Golden Gate assembly methods. For vectors containing multiple activation domains (ADs), the ADs were separated by short glycine-serine linkers. All plasmids used are listed in, Table 1 and were privately deposited to Addgene and will be publicly released (respective, anticipated Addgene IDs can be found in Table 1). All Sa-dCas9 plasmids were based on nuclease-null *Staph aureus* ortholog as described in Ran et al. 2015 as generously donated by the Zhang lab, ST1-dCas9 plasmids were based on M-ST1n-VP64 (Addgene #48675). All gRNAs were selected to bind anywhere between 1 and 500 base pairs upstream of the transcriptional start site. All gRNAs were expressed from either cloned plasmids (Addgene #41817) or cloned into single vector activator plasmids. Reporter targeting gRNAs were previously described (Addgene #48671 and #48672).

**Table 1.**
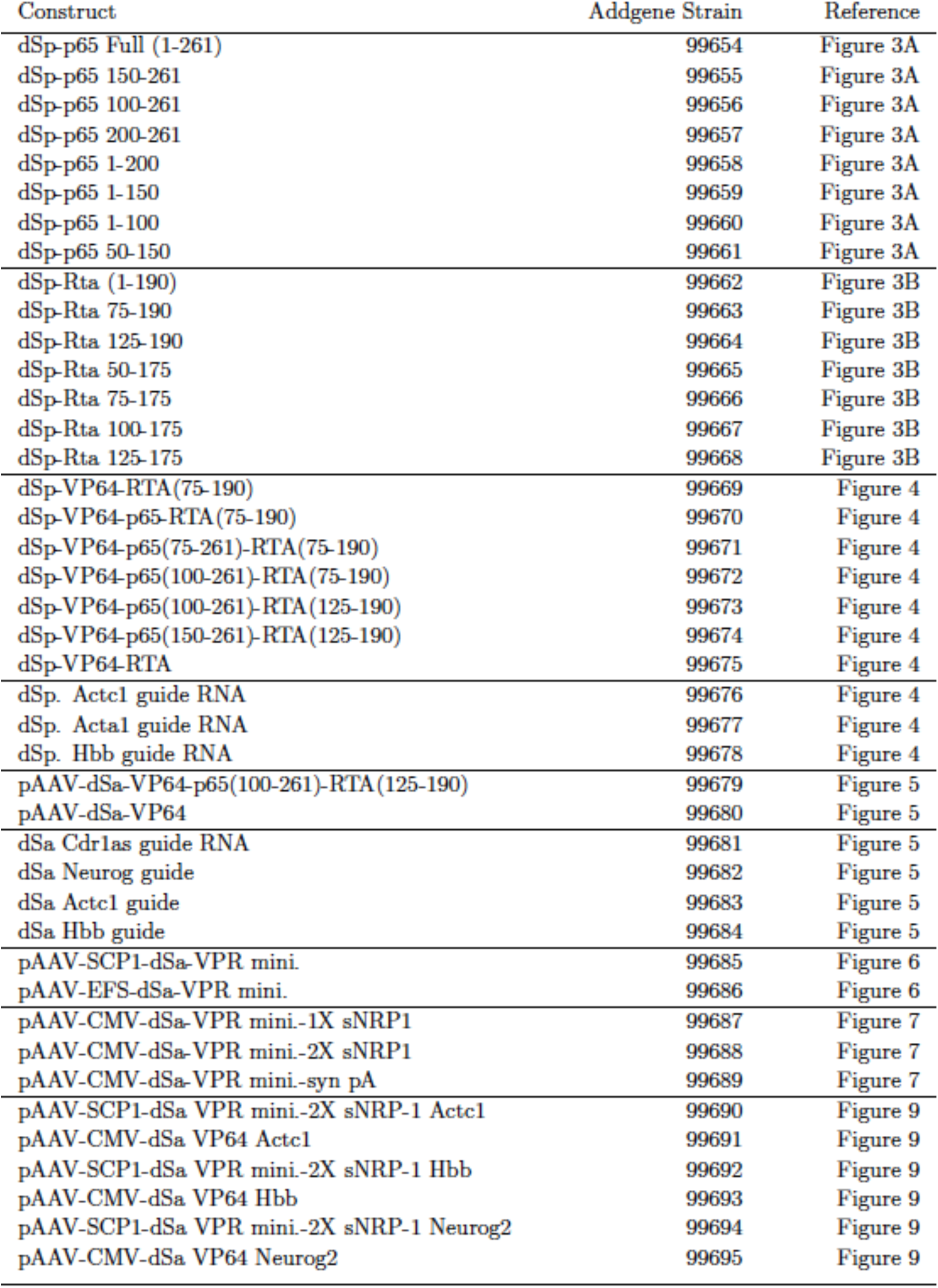
Reference of all constructs designed, and deposited to Addgene, along with respective Addgene strain IDs.

| Construct | Addgene Strain | Reference |
| --- | --- | --- |
| dSp-p65 Full (1-261) | 99654 | Figure 3A |
| dSp-p65 150-261 | 99655 | Figure 3A |
| dSp-p65 100-261 | 99656 | Figure 3A |
| dSp-p65 200-261 | 99657 | Figure 3A |
| dSp-p65 1-200 | 99658 | Figure 3A |
| dSp-p65 1-150 | 99659 | Figure 3A |
| dSp-p65 1-100 | 99660 | Figure 3A |
| dSp-p65 50-150 | 99661 | Figure 3A |
| dSp-Rta (1-190) | 99662 | Figure 3B |
| dSp-Rta 75-190 | 99663 | Figure 3B |
| dSp-Rta 125-190 | 99664 | Figure 3B |
| dSp-Rta 50-175 | 99665 | Figure 3B |
| dSp-Rta 75-175 | 99666 | Figure 3B |
| dSp-Rta 100-175 | 99667 | Figure 3B |
| dSp-Rta 125-175 | 99668 | Figure 3B |
| dSp-VP64-RTA(75-190) | 99669 | Figure 4 |
| dSp-VP64-p65-RTA(75-190) | 99670 | Figure 4 |
| dSp-VP64-p65(75-261)-RTA(75-190) | 99671 | Figure 4 |
| dSp-VP64-p65(100-261)-RTA(75-190) | 99672 | Figure 4 |
| dSp-VP64-p65(100-261)-RTA(125-190) | 99673 | Figure 4 |
| dSp-VP64-p65(150-261)-RTA(125-190) | 99674 | Figure 4 |
| dSp-VP64-RTA | 99675 | Figure 4 |
| dSp. Actc1 guide RNA | 99676 | Figure 4 |
| dSp. Acta1 guide RNA | 99677 | Figure 4 |
| dSp. Hbb guide RNA | 99678 | Figure 4 |
| pAAV-dSa-VP64-p65(100-261)-RTA(125-190) | 99679 | Figure 5 |
| pAAV-dSa-VP64 | 99680 | Figure 5 |
| dSa Cdr1as guide RNA | 99681 | Figure 5 |
| dSa Neurog guide | 99682 | Figure 5 |
| dSa Actc1 guide | 99683 | Figure 5 |
| dSa Hbb guide | 99684 | Figure 5 |
| pAAV-SCP1-dSa-VPR mini. | 99685 | Figure 6 |
| pAAV-EFS-dSa-VPR mini. | 99686 | Figure 6 |
| pAAV-CMV-dSa-VPR mini.-1X sNRP1 | 99687 | Figure 7 |
| pAAV-CMV-dSa-VPR mini.-2X sNRP1 | 99688 | Figure 7 |
| pAAV-CMV-dSa-VPR mini.-syn pA | 99689 | Figure 7 |
| pAAV-SCP1-dSa VPR mini.-2X sNRP-1 Actc1 | 99690 | Figure 9 |
| pAAV-CMV-dSa VP64 Actc1 | 99691 | Figure 9 |
| pAAV-SCP1-dSa VPR mini.-2X sNRP-1 Hbb | 99692 | Figure 9 |
| pAAV-CMV-dSa VP64 Hbb | 99693 | Figure 9 |
| pAAV-SCP1-dSa VPR mini.-2X sNRP-1 Neurog2 | 99694 | Figure 9 |
| pAAV-CMV-dSa VP64 Neurog2 | 99695 | Figure 9 |

### Cell culture and transfections

Neuro 2A, Hepa 1-6, and GC-1 cells were maintained in DMEM (High Glucose) with 4.5g/L D-Glucose, L-Glutamine, Sodium Pyruvate, and 10% supplemented heat inactivated FBS. C2C12 cells were maintained in the same medium, without Sodium Pyruvate. Prior to transfection, all lines were seeded in 24 well plates. The next day, cells were 40-50% confluent, and were transfected with either 200ng activator with 10ng of gRNA, or 200ng of single vector activator if the guide was included in the vector. All transfections were carried out with Lipofectamine 3000, using the standard protocol, with 1.5 uL of Lipofectamine 3000 and 2uL P3000 reagent per transfected well.

### qRT-PCR analysis

RNA was extracted from cells using the RNeasy PLUS mini kit (Qiagen) according to manufacturer’s protocol. 500ng of RNA was used to generate cDNA with the iScript cDNA synthesis Kit (BioRad), and 0.5ul of cDNA was used for each qPCR reaction, utilizing the KAPA SYBR® FAST Universal 2X qPCR Master Mix.

### Large Scale AAV-DJ production and purification

Large-scale polyethylenimine transfections of AAV cis, AAV trans, and adenovirus helper plasmid were performed in a ten-layer hyperflask (Corning) with near- confluent monolayers of HEK293 cells. Plasmids were transfected at a ratio of 2:1:1 (260 mg of adenovirus helper plasmid/130 mg of cis plasmid/ 130 mg of trans plasmid). PEI Max (Polysciences)/DNA ratio was maintained at 1.375:1 (w/w). After 60 hours, cells and supernatant were collected and freeze thawed 3 times alternating between a dry ice ethanol bath and 37°C incubator, spun down at 14,000g to sediment cell debris. The clarified feedstock was then concentrated by Tangential Flow Filtration down to roughly 15 mL. Prior to further concentration, ~20-30mL PBS was flown through large Amicon Ultra-15 (50 kDa MWCO) (Millipore) filters to wash out glycerin. Virus containing supernatant was poured into Amicon filters, with ~2mm space at top of filter (to prevent spilling of virus containing media out of filter, during centrifugation). Virus was spun at 3500g for 2 hrs, with a visual check at 1 hour mark to identify reduction in volume and ensure membranes did not dry out. The collected AAV supernatant was first treated with 50U/ml Benzonase (stock at 250U/uL) and 0.5U/ml Riboshredder (stock at 1U/uL) for 1 hour at 37 °C., shaking at 200-250 RPM. After incubation, the lysate was concentrated to <3 ml by ultrafiltration with Amicon Ultra-15 (50 kDa MWCO) (Millipore), and loaded on top of a discontinuous density gradient consisting of 2 ml each of 15%, 25%, 40%, 60% Optiprep (Sigma-Aldrich) in an 11.2 ml Optiseal polypropylene tube (Beckman-Coulter). The tubes were ultracentrifuged at 58000 rpm, at 18 °C, for 1.5 hr, on an NVT65 rotor. The 40% fraction was extracted, and dialyzed with 1xPBS (pH 7.2) supplemented with 35 mM NaCl, using Amicon Ultra- 15 (50 kDa or 100 kDa MWCO) (Millipore). The purified AAVs were quantified for viral titers, and stored in −80 °C.

### Small Scale AAV-DJ production and purification

Small scale AAV prep was initially performed by the Gene Transfer Vector Core of Schepens Eye Research Institute and Massachusetts Eye and Ear Infirmary. HEK293 cells were seeded in a 15cm dish, one day prior to transfection, such that cells reached 90% confluence in one day. The next day (day 2) adenovirus helper plasmid, pAAV cis plasmid, and pAAV DJ plasmid were transfected into cells by using polyethylenimine (PEI) at a ratio of 2:1:1 (26 µg of adenovirus helper plasmid, 13 µg of cis plasmid, and13 ug of trans plasmid, respectively). PEI Max (Polysciences, Warrington, PA)/DNA ratio was maintained at 1.375:1 (w/w). On day 5, cells and supernatant were collected, and subjected to 3 freeze-thaw cycles. The viral solution was spun down at 14,000 g at 4 C for 15 min. Then the supernatant was collected to new tubes. Tubes were transported on ice to the Wyss Institute for Biologically Inspired Engineering. Viral supernatants were spun down in Amicon Ultra-15 (50 kDa MWCO) (Millipore) filters for 40 min at 5000g in a fixed angle centrifuge. Viruses were stored at 4C overnight, each at roughly 2mL. The next day, 1uL Benzonase (Millipore, 101695) was added to each virus, resulting in >= 125U Benzonase per 1 mL virus. Each tube was rocked back and forth on bench top, at room temperature and placed in 37 C room for 30 min. for Benzonase treatment. Treated virus was spun down at 4800g on a tabletop centrifuge for 15 min. at 4 °C to sediment cell debris. Buffer exchange was performed twice with Amicon Ultra-15 (50 kDa MWCO) (Millipore) filters, bringing volume up to 15mL with PBS (-CaCl, - MgCl2) + 35 mM NaCl. For concentration, buffered virus was spun down in Amicon filters for 30 min. at 5000g, at 4 °C to bring volume down to roughly 1mL. The purified AAVs were quantified for viral titers and stored at −80 °C.

### AAV-DJ titration via RT-qPCR

DNAseI treated samples using 2uL AAV in 20uL total rxn. with 1uL enzyme at 37C for 30 min. Standard qPCR was run on sample alongside dilution series of previously linearized template vector; titer was determined to be approximately 1e11.5 vg/mL for small scale preps used for transduction.

### AAV transduction in cell lines

Transduced N2As, GC-1, C2C12, and Hepa 1-6 seeded in 24 well plate with 10uL each vector. Mixed 10uL virus with 100uL media (in master mixes) and laid that down on the cells, then dispensed additional 500uL cell line specific media on top. Cell lines were untouched for 3 days, media switched, and then media switched every 2 days until RNA collection on day 9.

